# The SARS-CoV-2 nucleoprotein associates with anionic lipid membranes

**DOI:** 10.1101/2023.09.15.557899

**Authors:** Mandira Dutta, Yuan Su, Gregory A. Voth, Robert V. Stahelin

## Abstract

Severe acute respiratory syndrome coronavirus 2 (SARS-CoV-2) is a lipid-enveloped virus that acquires its lipid bilayer from the host cell it infects. SARS-CoV-2 can spread from cell to cell or from patient to patient by undergoing assembly and budding to form new virions. The assembly and budding of SARS-CoV-2 is mediated by several structural proteins known as envelope (E), membrane (M), nucleoprotein (N) and spike (S), which can form virus-like particles (VLPs) when co-expressed in mammalian cells. Assembly and budding of SARS-CoV-2 from the host ER-Golgi intermediate compartment is a critical step in the virus acquiring its lipid bilayer. To date, little information is available on how SARS-CoV-2 assembles and forms new viral particles from host membranes. In this study, we find the N protein can strongly associate with anionic lipids including phosphoinositides and phosphatidylserine. Moreover, lipid binding is shown to occur in the N protein C-terminal domain, which is supported by extensive *in silico* analysis. Anionic lipid binding occurs for both the free and N oligomeric forms suggesting N can associate with membranes in the nucleocapsid form. Herein we present a lipid-dependent model based on *in vitro*, cellular and *in silico* data for the recruitment of N to M assembly sites in the lifecycle of SARS-CoV-2.

## Introduction

Respiratory viruses can easily spread from one person to another, hence causing many life-threatening disorders and sometimes may lead to epidemics or even pandemics. Severe acute respiratory syndrome coronavirus 2 (SARS-CoV-2) was the causative agent for the COVID-19 pandemic and responsible for hundreds of millions of cases and millions of fatalities globally (Sharma, Tiwari et al. 2020, Tsiotas and Tselios 2022). Since the first emergence of COVID-19 in Wuhan, China, numerous variants have been reported including the UK (B.1.1.7), South Africa (B.1.351), Delta (B.1.617), and Omicron (B.1.1.529) (Aleem, Akbar Samad et al. 2023, Carabelli, Peacock et al. 2023). Despite effective vaccines for reducing the disease burden and severity, there are limited small molecule treatment options (Akinosoglou, Schinas et al. 2022). To broaden the effectiveness of the SARS-CoV-2 inhibitors and potentially reduce the severity of the disease, it is crucial to improve our knowledge of the SARS-CoV-2 virus life cycle.

SARS-CoV-2 is a lipid-enveloped, single-stranded RNA virus with ∼30 kilobase genomic size(Brant, Tian et al. 2021). It encodes four major structural proteins, spike (S), membrane (M), envelope (E), and nucleocapsid (N) (Bangaru, Ozorowski et al. 2020, Zhang, Xiao et al. 2021, Jackson, Farzan et al. 2022). S protein is exposed on the surface of the virion and interacts directly with the receptor, angiotensin-converting enzyme 2 (ACE-2) during viral host cell entry (Hoffmann, Kleine-Weber et al. 2020), making it a prime target for drug design. The N protein is highly abundant in SARS-CoV-2 virions and in infected cells and has not been fully explored as a potential target for drug development against SARS-CoV-2 (Lang, Chen et al. 2021, Royster, Ren et al. 2023). N protein plays crucial roles in the virus life cycle including viral RNA replication and transcription (Baric, Nelson et al. 1988, Almazán, Galán et al. 2004). Viral genomic RNA and N protein assemble into a ribonucleoprotein (RNP) complex (Melchjorsen, Jensen et al. 2005, Hoffmann, Kleine-Weber et al. 2020), which interacts with M protein to package into virions (Narayanan, Maeda et al. 2000). To accomplish these tasks, N protein possesses an architecture consisting of an N-terminal RNA binding domain (NTD)(Khan, Hussain et al. 2022, Royster, Ren et al. 2023) and a C-terminal dimerization domain (CTD)(Baric, Nelson et al. 1988), and they are linked by an intrinsically disordered serine and arginine-rich linker. N protein is predominantly dimeric in solution (Chang, Sue et al. 2005, Zhou, Zeng et al. 2020) and can oligomerize through the CTDs (Chen, Chang et al. 2007, Ye, West et al. 2020) which are believed to bind the RNA packing signals during viral particle assembly (Luo, Chen et al. 2006). N also undergoes liquid-liquid phase separation in the phosphorylated form, which is proposed to regulate its function in viral genome processing (Carlson, Asfaha et al. 2020).

Despite knowledge on N protein interactions with RNA and higher order oligomerization, how N is recruited to sites of assembly for SARS-CoV-2 and other coronaviruses is not well understood. The membrane (M) protein is expressed in the Golgi and has been shown to bind N (Narayanan, Maeda et al. 2000, He, Leeson et al. 2004, Luo, Wu et al. 2006) and recruit N to assembly sites (Scherer, Mascheroni et al. 2022) likely via the C-terminal tail of M (Luo, Wu et al. 2006). While M alone doesn’t support virus-like particle (VLP) generation, co-expression of M with N can lead to VLP formation (Huang, Yang et al. 2004, Nguyen, Nguyen et al. 2022) with M, N, and E being most efficient for SARS-CoV-2 (Gourdelier, Swain et al. 2022, Nguyen, Nguyen et al. 2022). Nearly thirty years ago a murine coronavirus N protein was shown to associate with lipid membranes in both the free or nucleocapsid form (Anderson and Wong 1993, Wong and Anderson 1993). N binding to membranes was shown to be independent of the presence of other proteins and was driven predominantly by anionic membrane content (Anderson and Wong 1993, Wong and Anderson 1993). During SARS-CoV-2 infection, N has been detected at fused and convoluted membrane compartments, accumulating at regions that fold around ER membranes, a likely viral replication organelle (Scherer, Mascheroni et al. 2022). The role of host lipids in membrane localization of N and N incorporation into virions or VLPs has been underexplored. Many lipid-enveloped viruses that assemble and bud from the plasma membranes of host cells utilize lipid-protein interactions with a viral matrix protein (Freed 2006, Dick, Datta et al. 2015, Bobone, Hilsch et al. 2017, Favard, Chojnacki et al. 2019, Husby and Stahelin 2021, Motsa and Stahelin 2021, Norris, Husby et al. 2022) to mediate formation of viral progeny. Thus, in this study we examined the ability of the SARS-CoV-2 N protein to interact with lipid membranes *in vitro*, in cells and *in silico* and we performed molecular dynamics simulations to quantify the specific N/lipid interactions. Results shed light on the ability of N to interact with host cell lipids and recruitment of N to sites of SARS-CoV-2 assembly.

## Results and discussion

### The SARS-CoV-2 nucleoprotein associates with membranes containing anionic lipids

To test the hypothesis that the coronavirus N protein associates with lipid membranes, we first employed a lipid overlay assay (Shirey, Scott et al. 2017) using commercially available lipid strips (PIP strips and membrane lipid strips) containing an array of phosphoinositides, glycerophospholipids and several sphingolipids. The purified N protein from *E. Coli* was shown to associate with several anionic lipids at both room temperature and 4ºC (Figure 1A). Notably, N associated with all seven phosphoinositides (these include: PI(3)P, PI(4)P, PI(5)P, PI(3,4)P_2_, PI(4,5)P_2_, PI(3,5)P_2_, and PI(3,4,5)P_3_) as well as phosphatidylserine (PS) and phosphatidic acid (PA). The association of N with many of these lipids was based on electrostatics as increasing concentrations of NaCl (300 and 500 mM) reduced the binding to these anionic lipids albeit with PA to a lesser extent. Purified N protein was also able to associate with phosphoinositides on a membrane lipid strip (Figure 1A bottom panel) as well as PA and PS as seen in the PIP strips. N protein also associated with cardiolipin, an anionic lipid typically found in mitochondria but didn’t show appreciable association with cholesterol, sphingomyelin, phosphatidylinositol, diacylglycerol, or triacylglycerol. As with the PIP strips, the association of N with anionic lipids on the membrane lipid strip was also predominantly driven by electrostatics as increasing concentrations of NaCl reduced binding to these anionic lipids (Figure 1A).

**Figure. 1.**
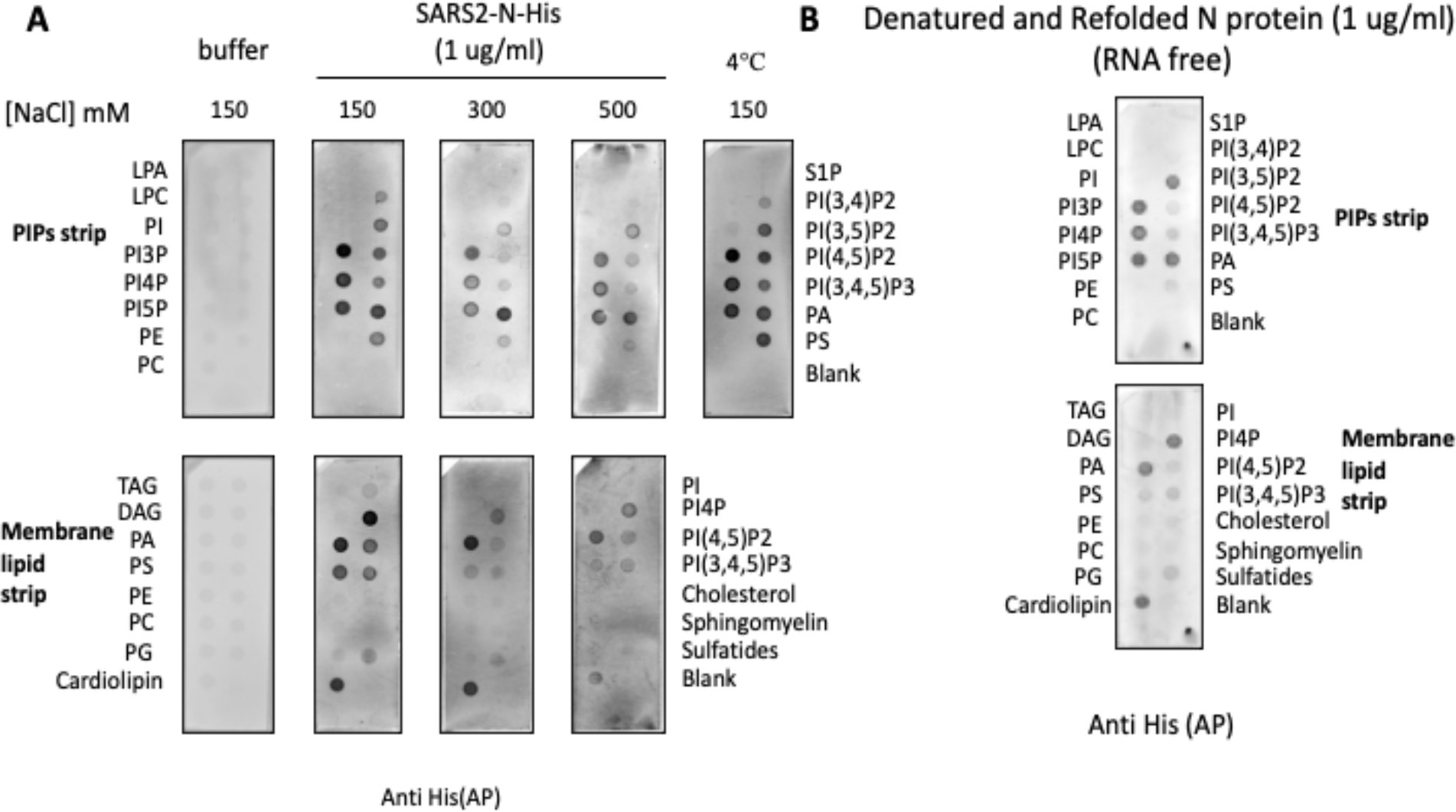
Lipid binding screen for the SARS-CoV-2 N protein. (A) Association of N protein with PIPs strip (top panel) and membrane lipid strips (bottom panel) reveal binding to anionic lipids including, phosphoinositides, PS, PA and cardiolipin. Increasing concentrations of NaCl reduces the binding of N protein to the anionic lipids (150 – 500 mM NaCl). (B) N binding with anionic lipid in the absence of RNA. Refolded N associates with anionic lipids on both the PIP and membrane lipid strips albeit to a lesser extent to PS, PI(4,5)P_2_, and PI(3,4,5)P_3_ than that of PA, PI(3)P, PI(4)P, PI(5)P, PI(3,5)P_2_ and cardiolipin.

The N protein is well known to bind RNA. While our N purification protocol contained enzymes to degrade RNA, we sought to confirm N binding to anionic lipids in the absence of RNA. Thus, we purified N using the same expression and purification protocol and then denatured and refolded N prior to lipid binding analysis. Refolded N protein was similarly able to associate with anionic lipids on both the PIP and membrane lipid strips (Figure 1B) albeit to a lesser extent to PS, PI(4,5)P_2_, and PI(3,4,5)P_3_ than that of PA, PI(3)P, PI(4)P, PI(5)P, PI(3,5)P_2_ and cardiolipin.

To confirm the lipid strip findings with membranes that more closely resemble the bilayers of biological membranes, we employed a large unilammelar vesicle (LUV) binding assay (Figure 2A). SARS-CoV-2 N exhibited strong association with anionic lipid containing vesicles (the seven phosphoinositides, PA and PS) in contrast to much lower association to vesicles containing only zwitterionic lipids (Figure 2B). In consonance with the electrostatic dependent association detected for N with lipid strips, N binding to anionic lipid vesicles was markedly reduced in the presence of 500 mM NaCl. A high throughput put surface plasmon resonance binding assay was also used to test binding of lipid vesicles to N (Figure 3). For these experiments, a Ni-NTA chip was used to capture N to take advantage of lipid vesicles of different composition injections on eight channels of the Biacore 8K instrument. Anionic lipid vesicles associated with N in a concentration dependent manner for the seven phosphoinositides as well as vesicles containing PA or PS (Figure 3).

**Figure 2.**
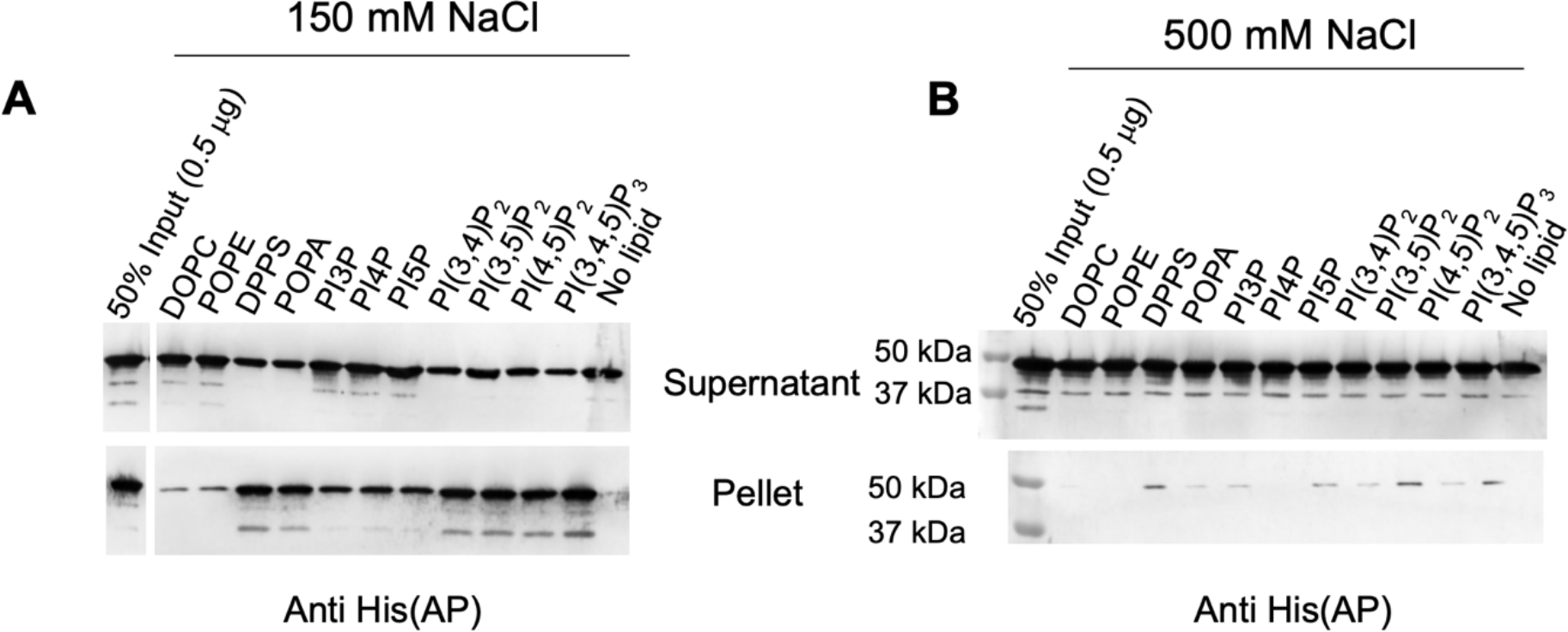
Large unilammelar vesicle binding assay for SARS-CoV-2 N and different anionic lipids. (A) Association of the N protein with anionic lipid containing vesicles in contrast to the vesicles containing only zwitterionic lipids. The top panel is the supernatant and the bottom panel the lipid pellet (protein bound fraction) for the centrifugation assay. (B) The vesicle binding assay was completed at 500 mM NaCl demonstrating a reduction in anionic lipid binding akin to the results in figure 1.

**Figure 3.**
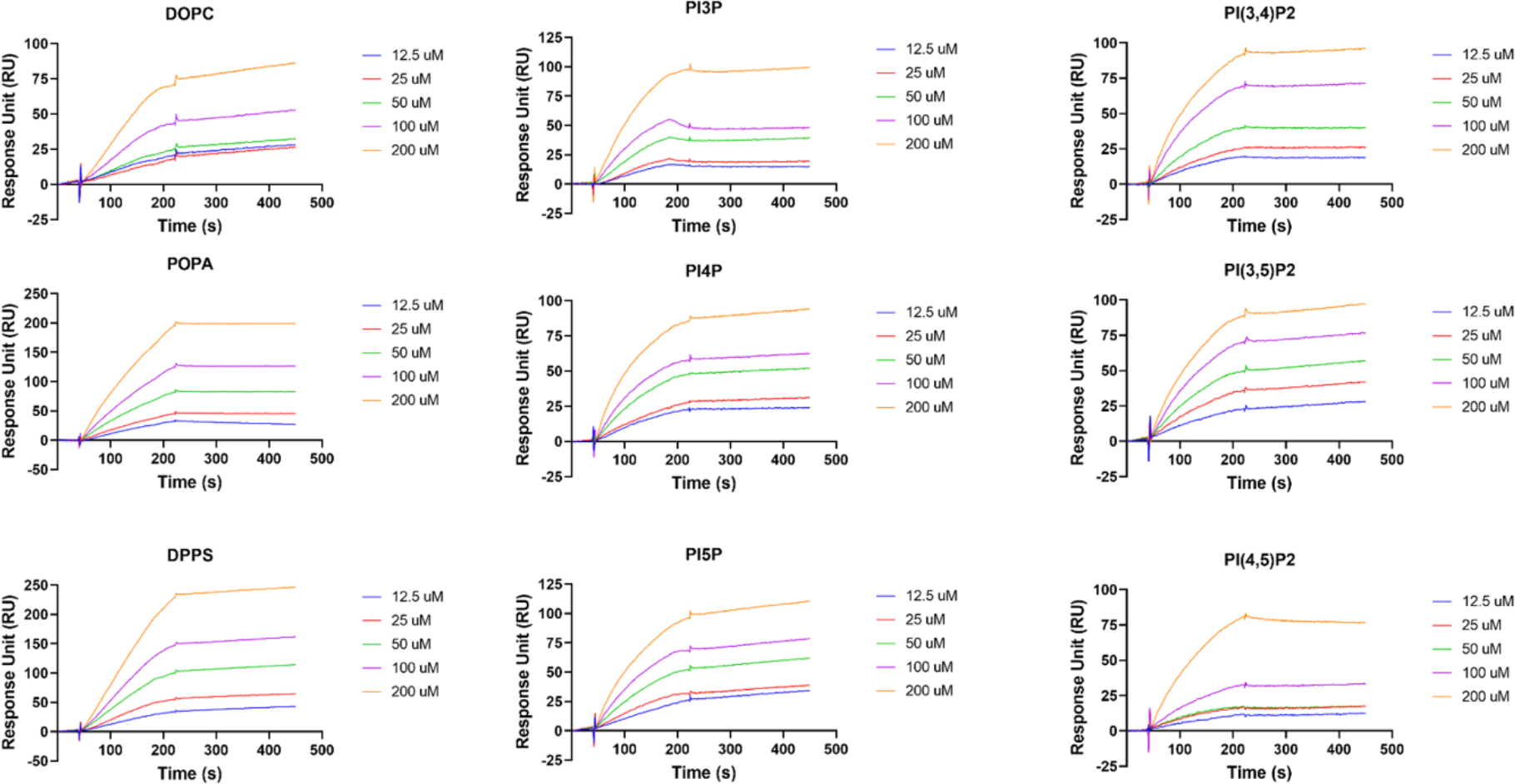
Surface plasmon resonance lipid vesicle binding assay for the SARS-CoV-2 N protein. Binding of lipid vesicles (anionic lipid labelled in each panel) to N in a high throughput surface plasmon resonance binding assay. Association of anionic lipid vesicles with N in a concentration dependent manner is shown for the seven phosphoinositides as well as vesicles containing PA or PS.

### The SARS-CoV-2 nucleoprotein can associate with anionic membranes in its oligomeric form and binds anionic membranes via its C-terminal domain

The N protein from SARS-CoV-2 and other betacoronaviruses can form oligomers (Chang, Chen et al. 2013, Carlson, Asfaha et al. 2020, Hsu, Chen et al. 2022), which play a critical role in assembly and binding RNA in SARS-CoV-2 genome packaging and virus spread. To determine if N oligomers can also associate with anionic lipid membranes, we crosslinked N using a BS^3^ cross linker (Figure 4A). Cross-linked N was able to associate with anionic lipids similar to purified N as shown in Figure 4B. This included association of N with the seven phosphoinositides as well as PA and PS. The association of cross-linked N with bis- and tri-phosphorylated PIPs was reduced compared to purified N somewhat akin to N that was denatured and refolded. Cross-linked N was also able to associate with lipid vesicles containing PA or PS as shown in Figure 4C confirming cross-linked N could associate with anionic lipids in the membrane bilayer environment.

**Figure 4.**
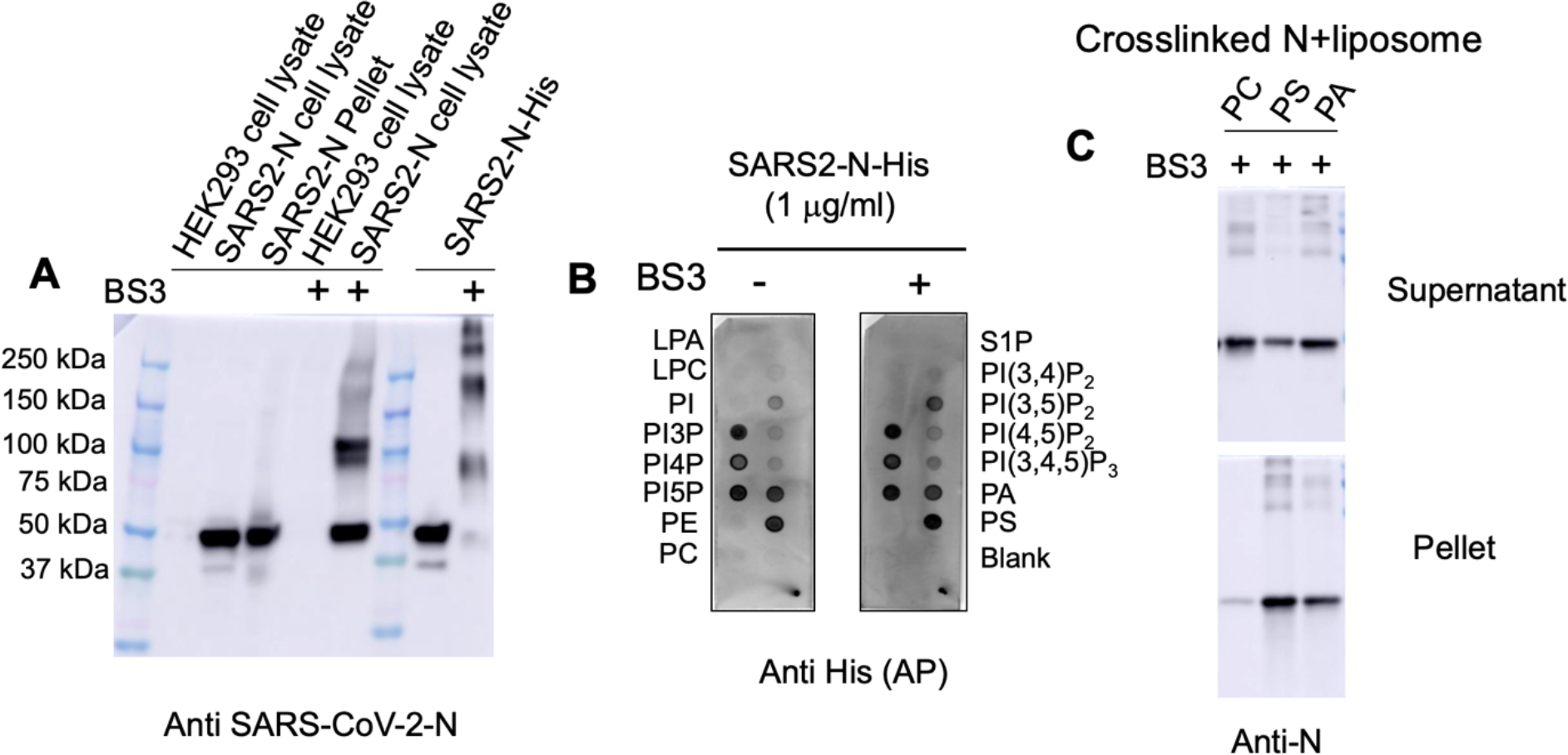
Association of N oligomers with anionic lipid membranes. (A) Western blot was performed on N protein expressed in HEK293 cells or from *E. Coli* with and without BS^3^ linker present demonstrating the formation of N protein oligomers. d(B) PIP strips were used to compare binding of the N protein in buffer and N protein post-crosslinking with BS^3^. N bound anionic lipids (particularly PI3P, PI4P, PI5P, PS and PA in the cross-linked or non-crosslinked form). (C) Cross-linked N was also able to associate with lipid vesicles containing PA or PS. The supernatant (top panel) and pellet (bottom panel) is shown for each lipid vesicle binding condition (PC, PS and PA).

To determine if a certain region of N associates with anionic membranes, we prepared several different N domain truncations as shown in Figure 5. This includes what we label as fragments 1-5 (F1-F5) where F1 corresponds to the N protein NTD (residues 1-181), F2 is the SR rich region and CTD (residues 182-419), F3 is the NTD and SR rich region (residues 1-247), F4 is the CTD alone (residues 248-419), and F is a fusion construct of the NTD and CTD (residues 1-181 and 248-419). As shown in Figure 5, F4 and F5 demonstrated lipid binding akin to full-length N albeit with lesser intensity of signal. F2 also displayed some detectable binding strongly suggesting N protein anionic lipid binding is attributed to its CTD due to binding by constructs F2, F4 and F5 (harboring the CTD) whereas F1 and F3 displayed little detectable binding where the CTD was lacking.

**Figure 5.**
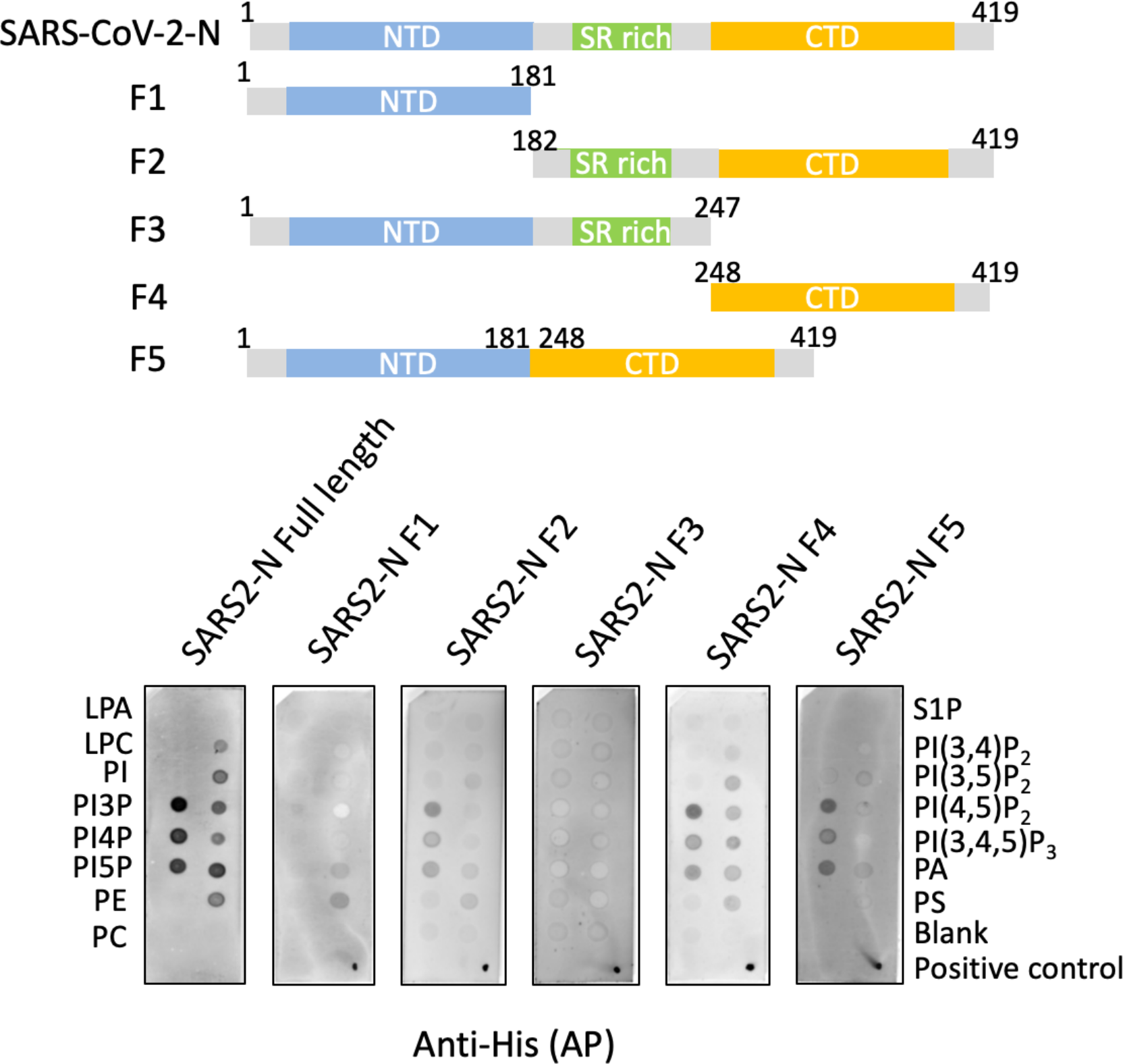
Association of different regions of N with anionic membranes. SARS-CoV-2 N and five different truncations of N were prepared (F1-F5) corresponding to the NTD (F1), SR rich region and CTD (F2), NTD and SR rich region (F3), CTD alone (F4), and a fusion construct of the NTD and CTD (F5), respectively.

### Computational Studies of the nucleoprotein C-terminal domain with lipid membranes

We next performed classical molecular dynamics (MD) simulations to investigate the protein-lipid interactions of N-CTD domains with different biologically relevant lipid membranes (see Table. 1 and Figure 6A-C). Here we re-numbered each CTD domain starting from 1 and named each monomer as CTD1 and CTD2.

**Table 1.**
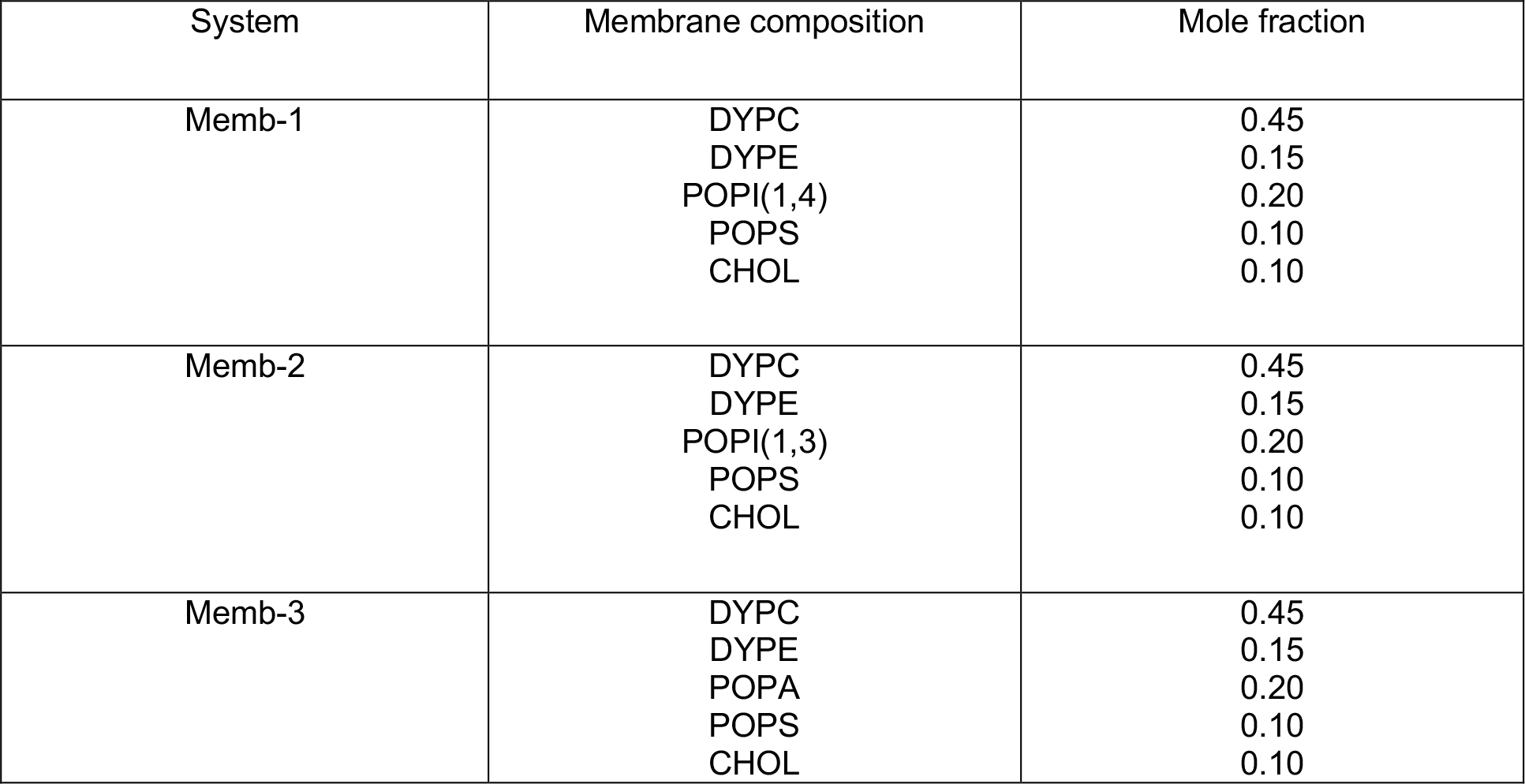
Simulation systems.

**Figure 6.**
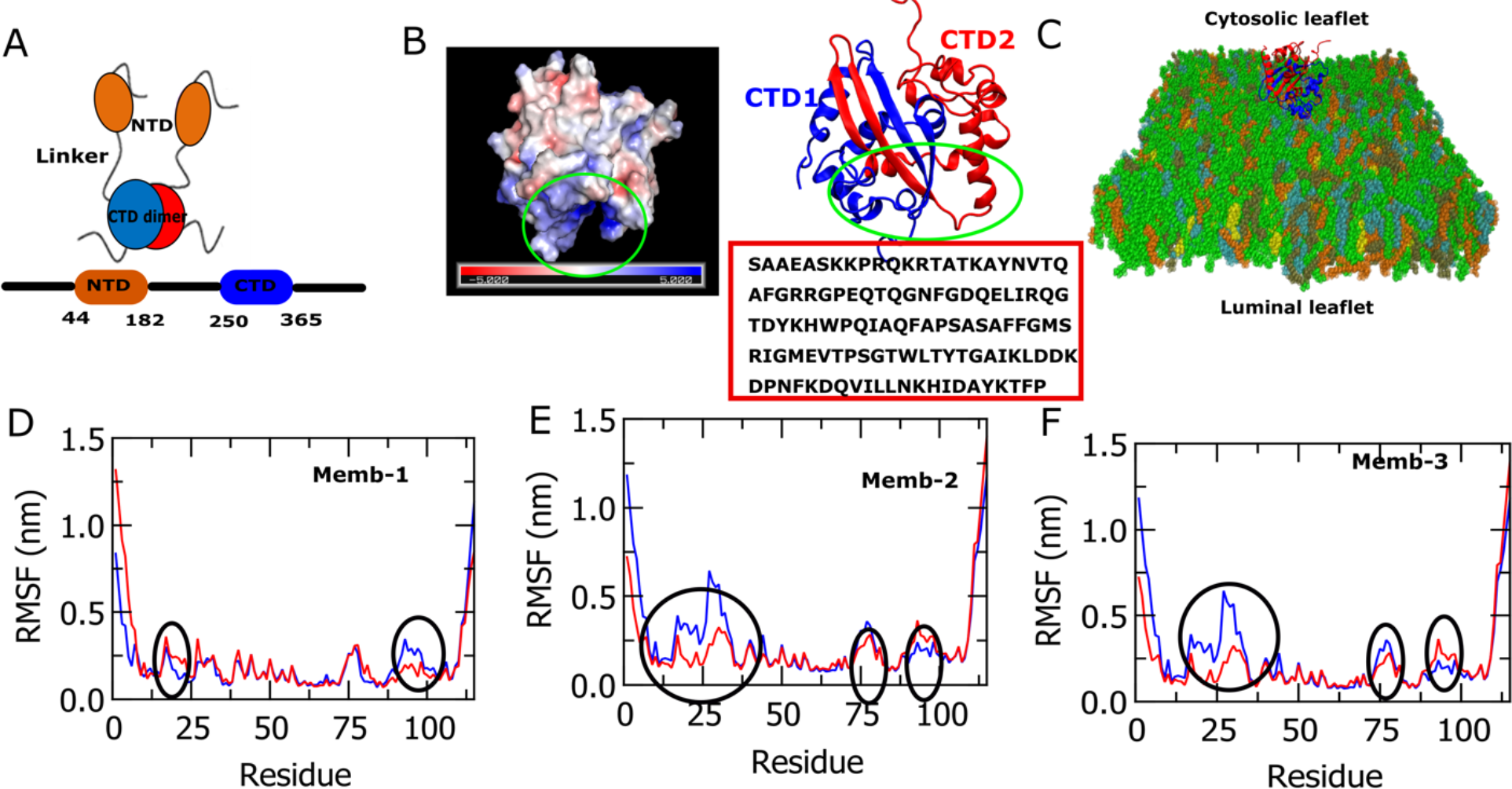
Structure of C-terminal domain (CTD) of SARS-CoV-2 N protein. Root means square fluctuations (RMSF) and electrostatic surface plots of N-CTDs. (A) Schematic diagram of N protein. NTD and CTD domains are connected via a linker region where the CTD domain forms a dimer. (B) The crystal structure of N protein was taken from PDB ID-7DE1. The electrostatic surface plot of N-CTDs shows the membrane binding surface has a high number of positively charged residues. Red box shows the amino-acid sequence of N-CTD domain. (C) N-CTD dimers bound with the lipid membrane. D, E, and F plots show RMSF of CTD1 and CTD2 domains. The black circles point out the regions where one monomer shows higher fluctuation value compared to the other monomer, referring one CTD shows significant interactions with the lipid membrane and the other is not.

We performed 1 μs MD production runs for three different membrane models with 2 replicas for each of them. Figures 6D, E, and F show the root means square fluctuation (RMSF) of CTD1 and CTD2 for memb-1, memb-2, and memb-3 models, respectively. The black circles point out the regions where one monomer shows lower fluctuation values than the other which indicates one monomer has significantly higher interactions with the lipid membrane than the other. The electrostatic surface plot (Figure 6B) shows that the binding interface is rich in positively charged residues. To understand the protein-protein and protein-lipid interactions in the different membrane models, we have calculated the number of contacts of CTD1-CTD2 and CTD dimer with lipids using a cut-off distance of 0.6 nm (Figure 7). For protein, Cα atoms were considered, and the heavy atoms were chosen for the lipid species. It can be seen that the memb-1 model shows a lower number of protein-protein contacts compared to memb-2 and memb-3 models (Figure 7A(i)). In contrast, for the protein-lipid contacts, it is observed that memb-1 shows a higher number of contact formations than the other two models (Figure 7A(ii)). Next, we examined the contacts of CTD dimers with PI and PA lipids. We noticed that the memb-1 shows significantly higher number of contacts between CTDs and PI(1,4) lipids than the contacts with PI(1,3) and PA lipids in memb-2 and memb-3, respectively, which indicates anionic PI(1,4) lipid has a significant contribution in overall lipid-protein interactions (Figure 7A(iii)). The contact analysis also signifies that for memb-1 N-CTD dimer forms robust protein-lipid interactions especially with anionic PI(1,4) lipids over protein-protein interactions. However, the memb-3 model shows an opposite behavior where N-CTD dimer does not significantly interact with PA lipids.

**Figure 7.**
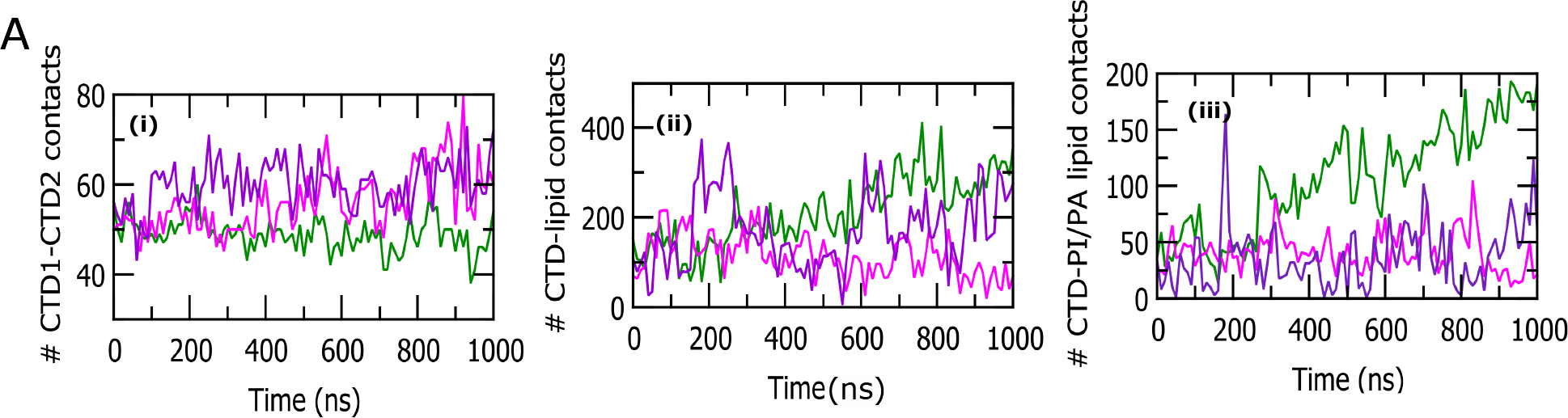
Number of contacts between N-CTD monomers and lipid species. (A) Number of contacts vs. simulation time. (i), (ii) and (iii) show number of interdomain contacts of CTD1 and CTD2, number of contacts between CTD dimers and lipid species, and number of contacts between CTD dimers and PI or PA lipids respectively. Green, magenta and violet colors indicate memb-1, memb-2 and memb-3. We consider 0.6 nm as a cut off distance for the calculation of contact pairs. For protein-protein contact we consider Cα atoms and for protein-lipid contacts we take Cα atoms of protein and heavy atoms of lipid species.

To analyze further, we have calculated the percentage occupancy of PI and PA lipids around N CTD residues (Figure 8). For the memb-1 model, we observe CTD1 shows higher occupancy of PI(1,4) lipids than the other monomer (CTD2) (Figure 8A(i) and (ii)). Here, we have picked out the residues which have occupancies greater than 50%. The residues with more than 50% occupancies are 1SER, 2ALA, 6SER, 7LYS, 10ARG, 13ARG, 17LYS, 22THR, 23GLN, 27ARG, 44ARG, and 112LYS (Figure 8D(i)).

**Figure 8.**
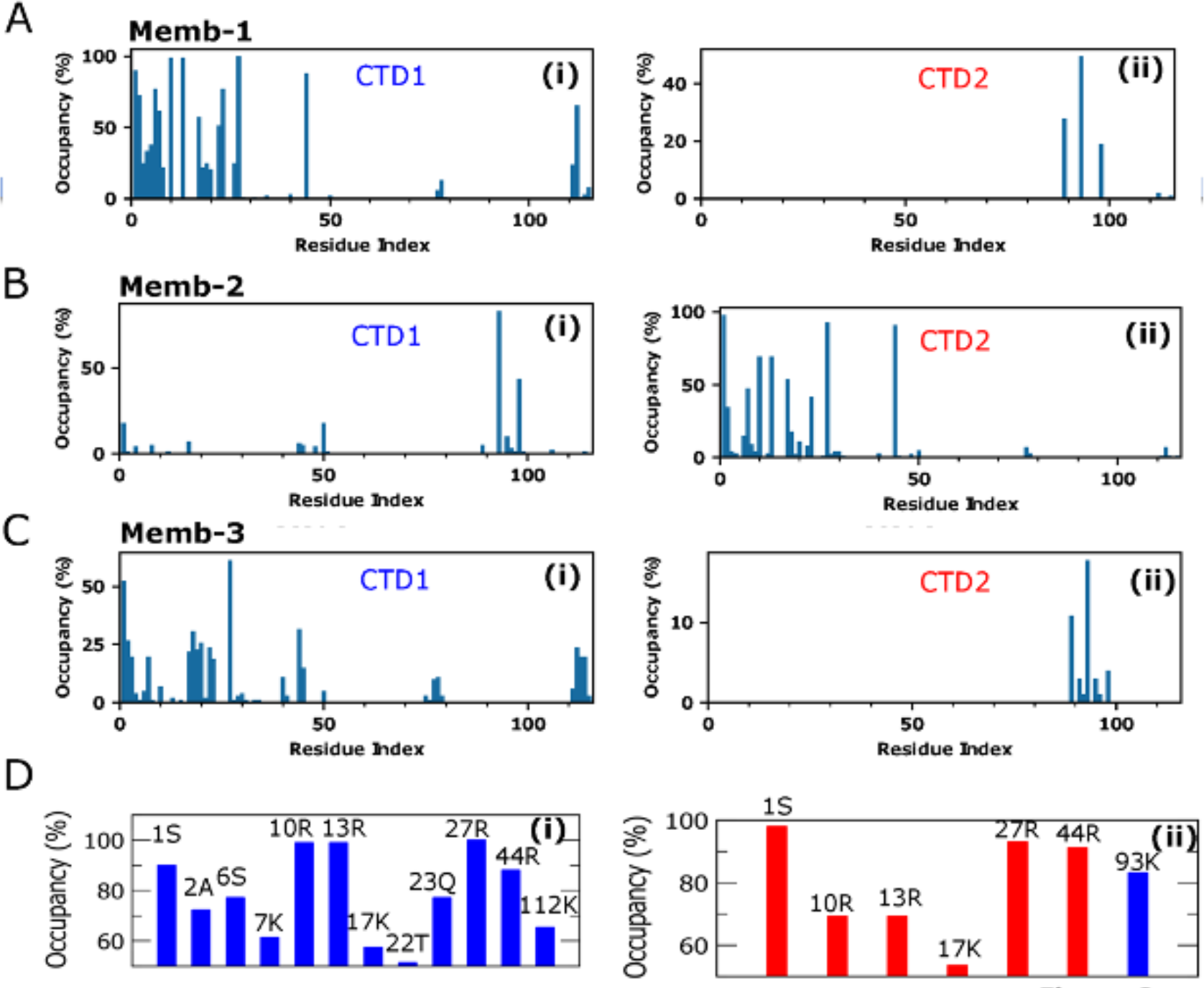
Percent of PI(1,4), PI(1,3) and PA lipid occupancies around protein residues. (A) (i) and (ii) show the % occupancy of PI(1,4) lipids vs. CTD1 and CTD2 residues in memb-1 model. (B) % occupancy of PI(1,3) lipids vs. CTD1(i) and CTD2 (ii) in memb-2 model. (C) % occupancy of PA lipids vs. CTD1 (i) and CTD2 (ii) in memb-3 model. (D) (i) Shows the residues which have occupancies of PI(1,4) lipids (memb-1) greater than 50%. (ii) Shows the residues which have occupancies of PI(1,3) lipids (memb-2) greater than 50%. Bule and red colors indicate CTD1 and CTD2, respectively.

Similarly, for memb-2 CTD2 shows more residues with higher occupancies than CTD1 (Figure 8B) and the residues with occupancies more than 50% are 1SER, 10ARG, 13ARG, 17LYS, 27ARG, 44ARG, and 93LYS (Figure 8D(ii)). The residues with higher occupancies are important for the interactions with the lipid species. For memb-3, a single residue (27ARG) is observed with an occupancy of more than 50% (Figure 8C). To understand the kinetics of lipid interactions at the protein surface, we have also calculated the residence time of interactions which quantifies the interaction time of each lipid head group at the protein-binding interface. Residence time provides insights into the dynamical behavior of the lipid species. Here, the calculations were done for PI and PA lipids of three different membrane models. We observe a diverse kinetic profile for protein-lipid interactions for memb-1, memb-2, and memb-3 models (Figure 9A, B, C). We identified the residues which have residence times greater than 400 ns (Figure 9 A(iii), B(iii)). The residues are 6SER, 7LYS, 10ARG, 13ARG, 19TYR, 22THR, 23GLN, 26GLY and 27ARG for PI(1,4) (memb-1) and 10ARG, 13ARG, 22THR, 27ARG, and 44ARG for PI(1,3) (memb-2). Figure 9D shows representative structures for protein-lipid interactions for memb-1 and memb-2 models. A comparison of Figure 8 and Figure 9 helps to quantify the common residues with higher PI lipid occupancies and higher residence time of interactions. It provides useful insight about the CTD residues which play an important role in protein-lipid interactions. Figure S1 shows lipid counts of PI and PA lipids around protein residues which refers to the distribution of specific lipids around protein residues. All these studies impart the idea of significantly different membrane responses depending on the different composition of the membrane structure.

**Figure 9.**
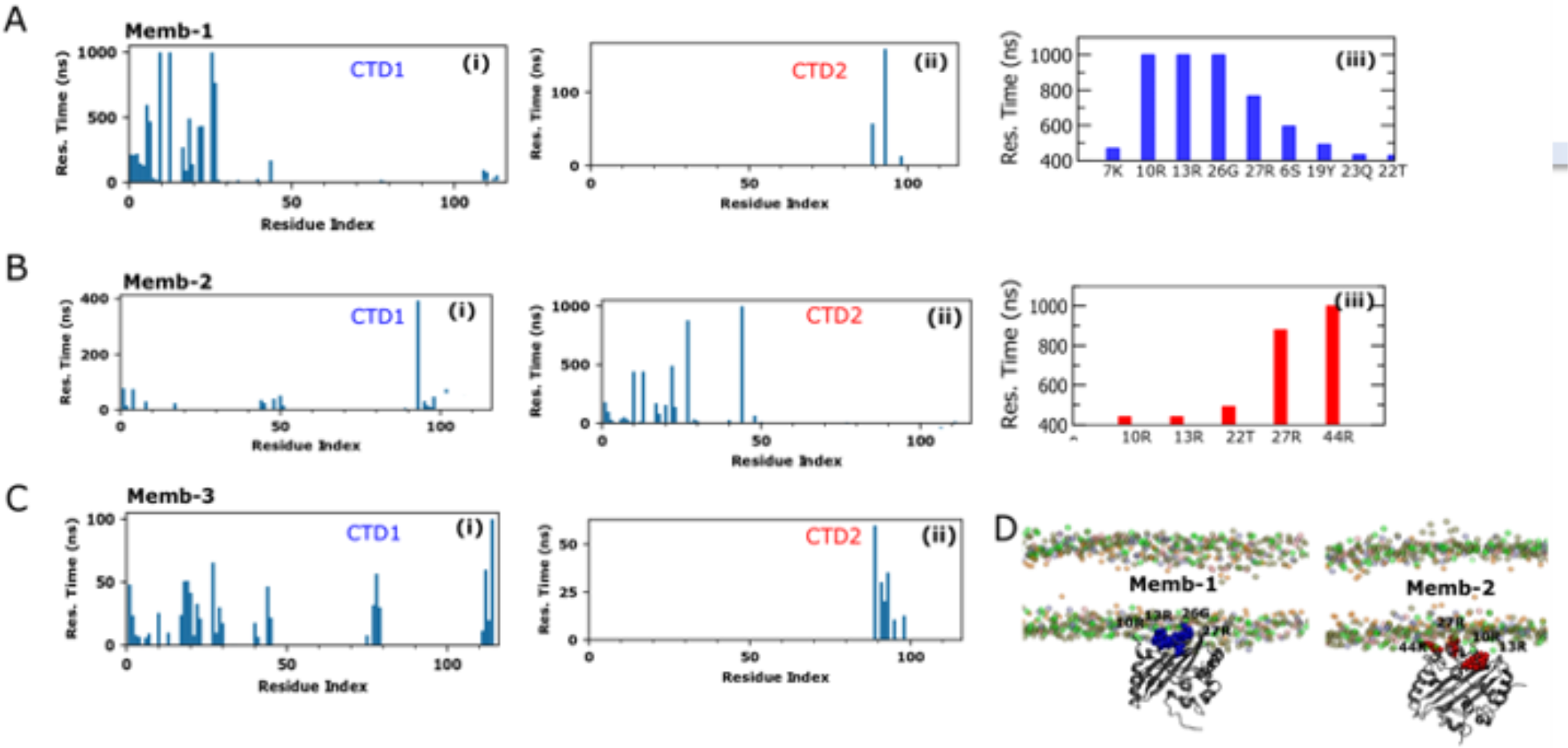
Residence time of interaction between PI(1,4), PI(1,3) and PA lipids, and protein residues. (A) (i) and (ii) show residence time of PI(1,4) lipid vs. CTD1 and CTD2 respectively for memb-1 model. (iii) Shows the residues which have residence times greater than 400 ns. (B) Residence time of interaction of PI(1,3) with CTD1 (i) and CTD2 (ii) for memb-2. (iii) Shows the residues which have residence time greater than 400 ns. (C) Residence time of interaction of PA with CTD1 (i) and CTD2 (ii) for memb-3. (D) Shows the representative structure of N-CTDs with lipid membranes for memb-1 and memb-2. Blue and red colors indicate CTD1 and CTD2 respectively. Green, pink, blue, tan and orange colors represent lipid head groups of PI or PA, PS, PE, PC and cholesterol, respectively.

To explore further, we have calculated the lipid density maps averaged over the last 50 ns of the MD simulation trajectories (Figure S2). The darker region indicates a higher density of specific lipids. We observe a higher accumulation of PI(1,4) lipids around the N-CTD dimer for memb-1 (Figure S2A). Memb-2 shows an accumulation of both PI(1,3) and PE lipids around protein (Figure S2B). However, for memb-3 we observe more accumulation of PS lipids compared to PA lipids (Figure S2C). Figure 10 shows the deformation pattern of membrane surface in different membrane models which indicates the location of each lipid phosphate group in the XY plane averaged over the last 50 ns of simulation trajectories. The red to blue color bar indicates gradual elevation of membrane surfaces along central axis towards the cytosolic leaflet. Memb-1 shows elevation of the membrane surface at the center towards cytosol. The CTD dimer shows higher interactions with lipid headgroups specifically anionic PI(1,4) lipids and pulls them towards the cytosol (Figure 10A). For memb-2 and memb-3 (Figure 10B, 10C), we observe slight elevation at the membrane corner. This result sheds light on the possible mechanisms to generate the curvature in the membrane surface which is an important phenomenon for the viral assembly and budding processes.

**Figure 10.**
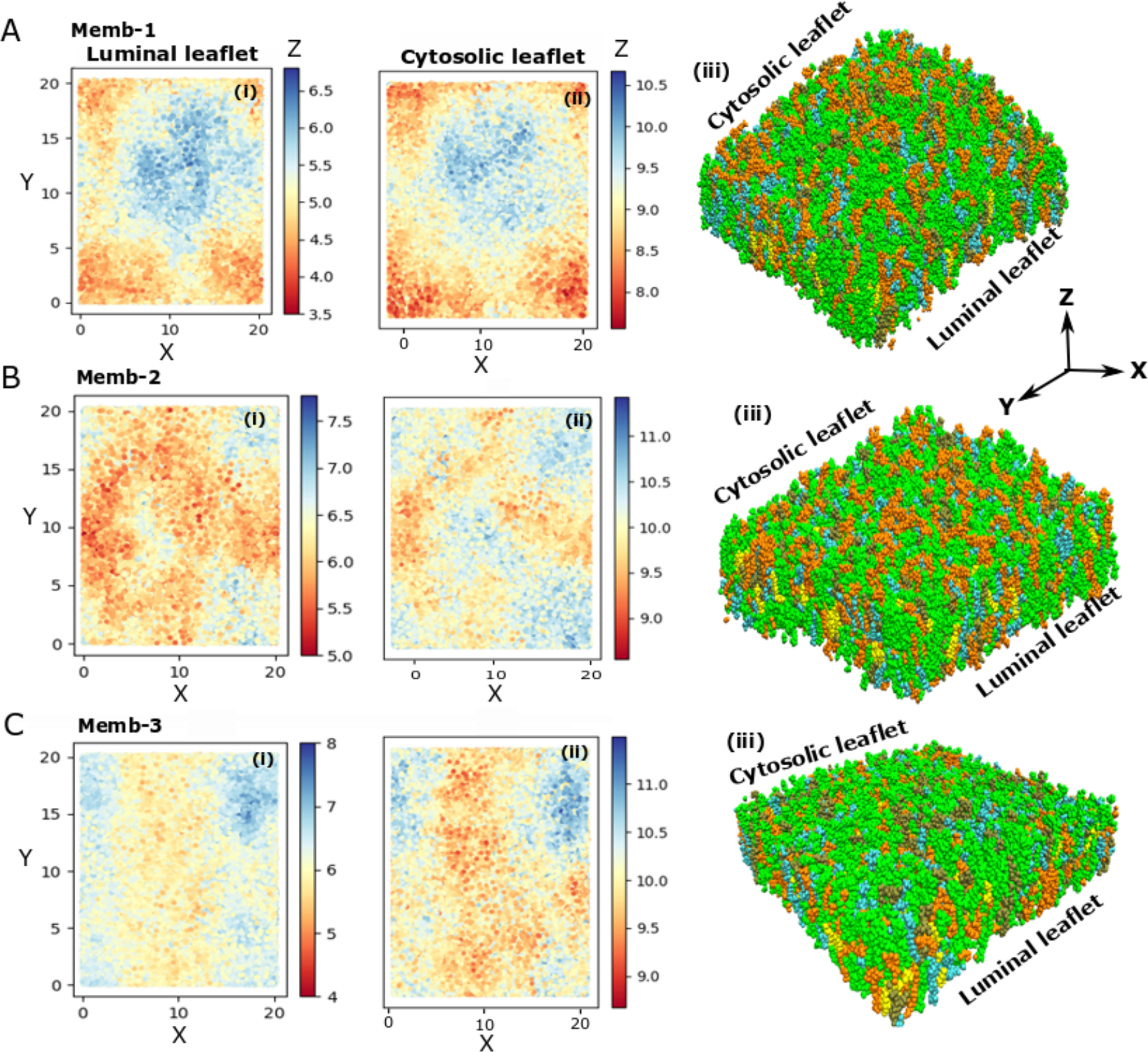
Membrane response due to N-CTD/lipid interactions. (i) and (ii) show the luminal and cytosolic leaflets for (A) memb-1, (B) memb-2 and (C) memb-3, respectively (iii) shows corresponding membrane representation. Each plot in (i) and (ii) shows average position of the lipid phosphate groups calculated over the last 50 ns of the MD trajectories. The color bar from red to blue refers to the gradual elevation of the membrane surface towards cytosolic leaflet.

## Conclusions

Our study provides important insights into the molecular interactions of the SARS-CoV-2 N CTD with lipid membranes. Both our experimental and simulation studies show association of N protein with anionic lipids. The association of N protein with the majority of the lipid components analyzed is based on lipid-protein electrostatic interactions. The electrostatic surface charge of the N protein CTD reveals a positively charged binding surface, which facilitates the interactions with anionic lipid components. We also observed that the N protein exhibits stronger association with anionic lipid containing vesicles compared to vesicles containing only zwitterionic lipids in a concentration dependent manner. To understand if N oligomers can also associate with anionic lipid membranes, we crosslinked N using a BS^3^ cross linker. Cross-linked N was able to associate with anionic lipids similar to purified N.

We were able to identify the protein residues important for the lipid-binding process from our MD simulations. Three membrane models of PI(1,4) (memb-1), PI(1,3) (memb-2) and PA lipids (memb-3) (Table.1) with dimeric N-CTD were used for the simulation studies. We notice that CTD in one monomer shows higher interactions with lipid membrane than the CTD in another monomer. Memb-1 exhibits lower protein/protein interactions and higher protein/lipid interactions compared to memb-2 and memb-3 models. Experiments with lipid strips and lipid vesicle sedimentation also shows that increasing concertation of NaCl reduces the association of N with PI(1,4) to a greater extent compared to PI(1,3). We identified specific residues of N-CTD with higher residence time of interaction with lipid head groups which corresponds to the stronger protein/lipid interactions. Notably, most of the residues with higher residence times are positively charged. Furthermore, we have quantified the lipid sorting pattern around CTD dimers and explored the nature of membrane deformation which may enhance the curvature formation for viral assembly and budding processes.

This work summarizes the specific N-protein/lipid interactions that may be critical for SARS-CoV-2 assembly and budding. How N protein is incorporated into virions or VLPs in the nucleocapsid form is not well understood, and our data indicates that in addition to M interactions with N, lipid interactions may play an essential role in assembly of N oligomers at the M residing membrane interface. Furthermore, N was recently found to reside at convoluted and fused membranes derived from the ER, a likely site of viral replication organelles. Thus, N-lipid interactions may be a key factor in localization of N to these sites where N plays an important role in transcription and genome replication. Understanding the mechanisms of N-lipid interactions is crucial for further understanding of the SARS-CoV-2 life cycle and could help spawn development of novel treatments against SARS-CoV-2 and related viruses.

## Materials and methods

### N protein expression and purification

A N protein expression plasmid (pET28a(+)) with a C-terminal His_6_ tag was codon optimized for *E. Coli* expression and generated by GeneUniversal (Newark, DE). Rosetta™ (DE3)pLysS cells (Millipore Sigma, Burlington, MA) were used for protein expression in LB media. Following an overnight starter culture of 10 mL, 2 L of LB media was inoculated and grown to O.D._600 nm_ ∼ 0.55. At this time, 0.1 mM (IPTG) was added to induce N protein expression and the cells were kept with shaking at 200 rpm at 37ºC for 3 hours. Cells were then collected using centrifugation at 6,000 *x g* at 4ºC and the N protein was purified by affinity chromatography using Ni-NTA agarose (Redwood City, CA) according to the manufacturer’s protocol. The N protein obtained from affinity chromatography was then further purified by size exclusion chromatography using a HiLoad 16/600 Superdex 200 pg column (Cytiva, Marlborough, MA). The resulting N fractions from size exclusion chromatography were concentrated to a final concentration of ∼ 1.5 μg/μl for each preparation.

### N-lipid interactions

PIP Strips™ and Membrane Lipid Strips™ were from Echelon Biosciences, Inc. (Salt Lake City, UT). N protein binding (1 μg/μL) to these strips was performed using an established protocol (Shirey, Scott et al. 2017) at room temperature and 150 mM NaCl unless noted.

Lipid vesicles were prepared as previously described (using 200 nm membranes for lipid extrusion) for a lipid pelleting assay to separate the bound and free forms of N via centrifugation (Johnson, Bhattarai et al. 2021). All lipids were from Avanti Polar Lipids, Inc. (Alabaster, AL) and used without further purification (1,2-dioleoyl-sn-glycero-3-phosphocholine (DOPC), 1-palmitoyl-2-oleoyl-sn-glycero-3-phosphoethanolamine (POPE), 1-palmitoyl-2-oleoyl-sn-glycero-3-phospho-L-serine (POPS), 1-palmitoyl-2-oleoyl-sn-glycero-3-phosphate (POPA), or 18:1 phosphoinositides). Control lipids (DOPC:POPE 70:30) were prepared and anionic lipids (10 mol% for POPS or POPA and 5 mol% for phosphoinositides) were added at the expense of DOPC for each lipid mixture. Total volumes for interaction analysis were 100 μL and N protein and lipid vesicles were incubated for 30 minutes at room temperature followed by centrifugation at 75,000 *x g* for 30 minutes to separate the lipid pound and free protein fractions. SDS-PAGE was then used to visualize the N protein in the supernatant and pellet fraction for each lipid vesicle composition tested.

SPR lipid binding assays were done using a Biacore 8K instrument and sensor chip NTA chip (Cytiva) for histidine tagged protein capture. In brief, the purified N protein harboring a His_6_ tag was captured on the surface of the NTA chip and then lipid vesicles of different compositions were flowed over the N protein surface at increasing concentrations. Lipid vesicles were prepared as previously described using extrusion through a 100 nm membrane (Adu-Gyamfi, Johnson et al. 2015).

### Cell culture maintenance and N protein expression

HEK293 cells were maintained at 37ºC and 5% CO_2_ in DMEM containing 10% FBS and 1% penicillin/streptomycin. The N protein plasmid pCAGGs-N (Plescia, David et al. 2021) was transfected using Lipofectamine™ 2000 (ThermoFisher Scientific, Waltham,

MA) and the N protein was allowed to express for 24 hours. The cells were then lysed, and the insoluble fraction separated by centrifugation. The supernatant was then used for cross-linking studies with BS^3^ (ThermoFisher Scientific) to demonstrate N protein oligomerization from cell lysates.

### Molecular dynamics simulations

In the present study, we choose three different membrane models where each model has five representative lipid species for the endoplasmic reticulum (ER) of eukaryotic cells(Monje-Galvan and Klauda 2015, Harayama and Riezman 2018, Monje-Galvan and Voth 2021, Dolan, Dutta et al. 2022). Each membrane model and the mole fraction of each lipid species are given in Table.1. For the N protein, we used the crystal structure of N-CTD dimer (PDB ID-7DE1) from the protein data bank(Yang, He et al. 2020). A 20 nm × 20 nm lipid bilayer for each membrane model was prepared using CHARMM-GUI Membrane Builder(Jo, Kim et al. 2007, Wu, Cheng et al. 2014, Lee, Cheng et al. 2016). Next, we calculated the electrostatic surface charge distribution of N-CTD dimers. We noticed accumulation of higher positive charge at a certain region of N-CTD (Figure 6B). Our experimental results show electrostatic interactions between anionic PI lipids and N-CTD. Based on the experimental observation, we placed the N-CTD dimer close to the center of the lipid bilayer such a way that the higher positive charge surface of CTD dimers bind with the membrane surface. We prepared two replicas for each membrane model with slight changes in the orientation of the CTD dimers on the membrane surface. We solvated each protein-lipid system in a solvation box of 20×20×14 nm^3^. To neutralize the system, we used 0.15M KCl salt. We performed MD simulations using an isobaric-isothermal ensemble on the GROMACS simulation package(Abraham, Murtola et al. 2015) with the CHARMM36m force field(Huang, Rauscher et al. 2017). We used Frontera (TACC), Anton2 (PSC), and Midway2 (RCC at the University of Chicago) machines to run simulations. A simulation time step of 2 fs and periodic boundary conditions were used for the simulations. 310.15 K temperature and a Nose-Hoover thermostat(Nosé 1984, Hoover 1985) were used with a coupling time constant of 1.0 ps in GROMACS. The pressure was set at 1bar with the Berendsen barostat(Berendsen, Postma et al. 1984) during initial relaxation. For the equilibration and production runs, we used the Parrinello-Rahman barostat (Parrinello and Rahman 1981) semi-isotropically with the compressibility of 4.5 × 10^−5^ and a coupling time constant of 5.0 ps. Non-bonded interactions were computed using a switching function between 1.0 and 1.2 nm, and for the long-range electrostatics Particle Mesh Ewald (PME)(Darden, York et al. 1993) was used. The LINCS algorithm(Hess, Bekker et al. 1997) was used to constrain hydrogen bonds. Each membrane model has two replicas and we performed 500ns equilibration and 1 μs production runs for each of them. The simulation trajectories were analyzed using internal GROMACS modules, and MDAnalysis(Michaud-Agrawal, Denning et al. 2011) and MDTraj(McGibbon, Beauchamp et al. 2015) python packages. We used Visual Molecular Dynamics (VMD) and PyMOL as visualizing software packages. Percent of lipid occupancy, residence time, and the number of lipids around proteins were calculated using PyLipID python package (Song, Corey et al. 2022).

## Acknowledgements

These studies were supported by the NIH (R01AI169896) to R.V.S and G.A.V. and the Purdue Pharmacy Live Cell Imaging Facility (NIH OD027034 to R.V.S.). Y.S. and R.V.S. are grateful for technical support by Dr. N. Dissinger and protein expression and purification support from Dr. D. Patra. M.D. thanks Dr. Viviana Monje-Galvan for insightful discussions on the simulation systems at the beginning of this work. The computations in this work were supported by the Frontera supercomputer at the Texas Advanced Computer Center (TACC), the Anton 2 machine at the Pittsburgh Supercomputing Center (PSC), and Midway2 at the Research Computing Center (RCC) of the University of Chicago.

