## Supporting information for "The SARS-CoV-2 nucleoprotein associates with anionic lipid membranes"

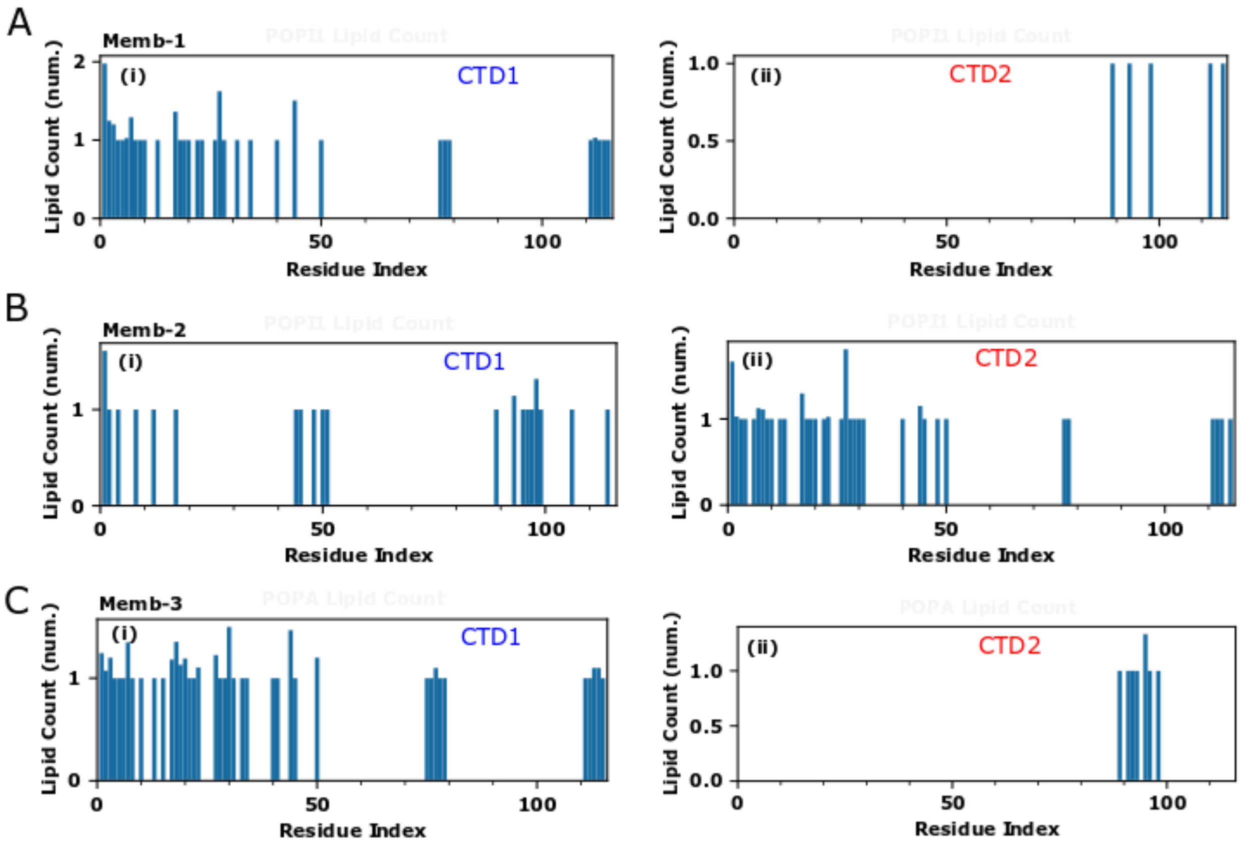


**Figure S1.** Number of PI(1,4), PI(1,3) and PA lipids around protein residues from MD simulations. (i) and (ii) correspond to the CTD1 and CTD2 and (A), (B), and (C) indicates PI(1,4) lipid for memb-1, PI(1,3) for memb-2 and PA for memb-3 models.

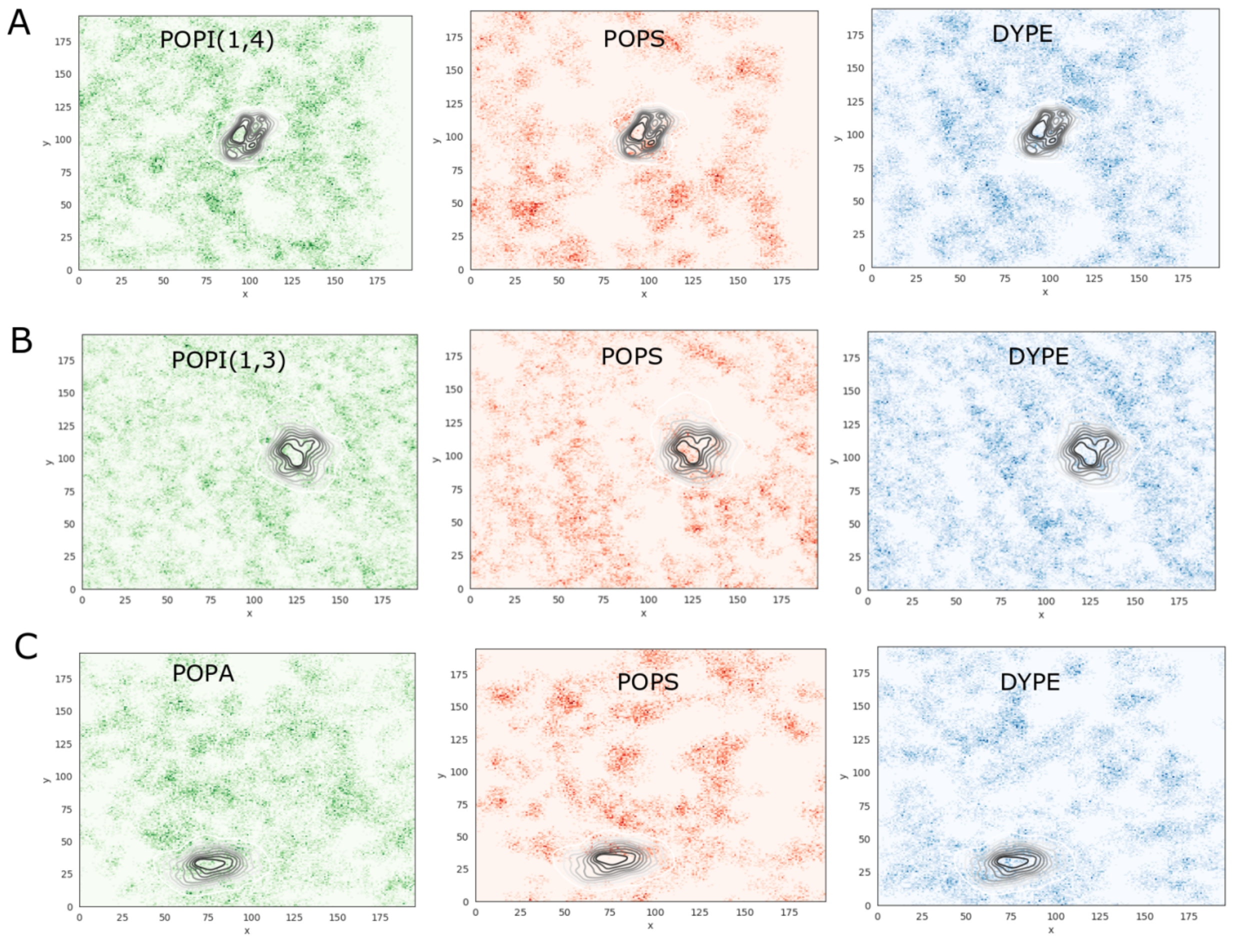


**Figure S2**. MD simulation lipid density maps relative to the location of the N-CTD dimers for (A) memb-1, (B) memb-2, and (C) memb-3 models. The darker regions indicate higher density of the respective lipid species.
